# Ketamine metabolism via hepatic CYP450 isoforms contributes to its sustained antidepressant actions

**DOI:** 10.1101/2024.04.03.587904

**Authors:** Thi Mai Loan Nguyen, Jean-Philippe Guilloux, Céline Defaix, Indira Mendez-David, Isabelle Etting, Jean-Claude Alvarez, Josephine C McGowan, Jaclyn N. Highland, Panos Zanos, Jacqueline Lovett, Ruin Moaddel, Emmanuelle Corruble, Denis J. David, Todd D. Gould, Christine A. Denny, Alain M. Gardier

**Author notes:** Corresponding author: Pr. Alain M. GARDIER Laboratoire de Neuropharmacologie Université Paris-Saclay, Inserm CESP/UMR 1018, Equipe MOODS, Faculté de Pharmacie, bâtiment HM1 recherche, bureau 4711, F-91400 Orsay, France.

## Abstract

(*R,S*)-ketamine (ketamine) has rapid and sustained antidepressant (AD) efficacy at sub-anesthetic doses in depressed patients. A metabolite of ketamine, including (*2R,6R*)-hydroxynorketamine ((*6*)-HNKs) has been reported to exert antidepressant actions in rodent model of anxiety/depression. To further understand the specific role of ketamine’s metabolism in the AD actions of the drug, we evaluated the effects of inhibiting hepatic cytochrome P450 enzymes on AD responses. We assessed whether pre-treatment with fluconazole (10 and 20 mg/kg, i.p.) 1 hour prior to ketamine or HNKs (10 mg/kg, i.p.) administration would alter behavioral and neurochemical actions of the drugs in male BALB/cJ mice with a highly anxious phenotype. Extracellular microdialysate levels of glutamate and GABA (Glu_ext_, GABA_ext_) were also measured in the medial prefrontal cortex (mPFC). Pre-treatment with fluconazole altered the pharmacokinetic profile of ketamine, by increasing both plasma and brain levels of ketamine and (*R*,*S*)-norketamine, while robustly reducing those of (*6*)-HNKs. At 24 hours post-injection (t24h), fluconazole prevented the sustained AD-like response of ketamine responses in the forced swim test and splash test, as well as the enhanced cortical GABA levels produced by ketamine. A single (*2R,6R*)-HNK administration selectively rescued the antidepressant-like activity of ketamine in mice pretreated with fluconazole within 24 hours of treatment. Overall, these findings are consistent with an essential role of (*6*)-HNK in mediating the sustained antidepressant-like effects of ketamine and suggest potential interactions between pharmacological CYPIs and ketamine during antidepressant treatment in patients.

## 1. Introduction

(*R,S*)-ketamine (ketamine) undergoes rapid and extensive metabolism by the hepatic cytochrome P450 (CYP) enzymes. Ketamine is first metabolized to *(R,S)*-norketamine, which is then further metabolized into 12 different *(R,S)*-hydroxynorketamines (HNKs) in humans, as well as in rodents (Can et al., 2016; Moaddel et al., 2015). Of these HNKs, (*6*)-HNK is the most abundant circulating HNK metabolite detected in the plasma of humans (Moaddel et al., 2015; Zarate et al., 2012; Zhao et al., 2012) and in the plasma and brains of rodents (Pham et al., 2018; Zanos et al., 2016).

Although the initial findings demonstrating antidepressant-relevant biological activity of (*2R,6R*)-HNK have been widely replicated and expanded upon by a number of independent research groups (Bonaventura et al., 2022; Cavalleri et al., 2018; Chen et al., 2018; Chou et al., 2018; Collo et al., 2018; Fred et al., 2019; Fukumoto et al., 2019; Pham et al., 2018; Wray et al., 2018; Yao et al., 2018; Ye et al., 2019), (2*R,*6*R*)-HNK is one of the major brain metabolites of (*R,S*)-ketamine, but its antidepressant-like activity and contribution to the actions of the parent drug has been contested (Abdallah, 2017; Collingridge et al., 2017; Zanos et al., 2017). We previously showed that a systemic or intra-cortical administration of (*6*)-HNK displayed similar sustained antidepressant-like behavioral actions to those of (*R,S*)-ketamine (i.e., at 24 hours post-injection (t24h)) when assessed using the forced swim test (FST). This effect was associated with enhanced glutamate and GABA release by pyramidal neurons and interneurons, respectively, in the medial prefrontal cortex (mPFC) (Pham et al., 2018). The results of some rodent studies suggest that (*2R,6R*)-HNK does not exert rapid and sustained antidepressant-like effects in different mouse preclinical tests of antidepressant effectiveness (Yamaguchi et al., 2018; Yang et al., 2017). It has also been reported that metabolism to (2*R,6R*)-HNK is not necessary for the antidepressant-like effects of the (*R*)-ketamine enantiomer (Yamaguchi et al., 2018). However, this conclusion contrasts with the finding that administration to mice of a 6-di-deuterated form of either ketamine or (*R*)-ketamine, which robustly hinder their metabolism to (*2S,6S*;2*R,*6*R*)-HNK and (*2R,6R*)-HNK respectively, attenuated the antidepressant-like behavioral actions in mice (Zanos et al., 2019; Zanos et al., 2016). Thus, the role of (*2R,6R*)-HNK in mediating ketamine’s antidepressant-like actions remains controversial. Further, the role of metabolism in the ketamine’s ability to enhance glutamate and GABA release in the mPFC has not been previously evaluated.

To address these questions, we assessed the behavioral and biochemical effects of (*R,S*)-ketamine administration following inhibition of hepatic CYP450 enzymes by fluconazole pretreatment (10 and 20 mg/kg, i.p.) in BALB/cJ and CD-1 mice. Fluconazole is a CYP450 inhibitor (CYPI) used as a strategy to study pharmacokinetic drug-drug interactions in humans (Peltoniemi et al., 2011). We hypothesized that if (*6*)-HNK metabolites are biologically active, fluconazole would reduce or prevent ketamine’s antidepressant-relevant behavioral and biochemical actions, coincident with decreasing brain exposure to these HNKs. By contrast, if metabolites are inactive, a CYPI would increase or not alter ketamine’s effectiveness in antidepressant-sensitive behavioral tests.

Similar to previous reports in mice (Humphrey et al., 1985; Santos et al., 2016) and rats (Uchida et al., 2014; Wang et al., 2017; Yang et al., 1996), we show here that fluconazole rapidly modifies the pharmacokinetic profile of (*R*,*S*)-ketamine in mice by increasing total brain exposure of (*R*,*S*)-ketamine and (*R*,*S*)-norketamine. Concomitantly, fluconazole robustly reduced (*2S,6S;2R,6R*)-HNK (*6*-HNK) levels. Notably, here, we show for the first time that fluconazole pre-treatment prevented the sustained antidepressant-like behavioral actions of ketamine at 24h following administration, and decreased ketamine-induced enhancement of cortical GABA release in the medial prefrontal cortex (mPFC) in BALB/cJ mice.

## 2. Materials and methods

### 2.1. Animals

For behavioral and microdialysis studies male BALB/cJRi (denoted as BALB/cJ hereafter) mice (7-8-weeks old) weighing 23-25 g at the beginning of the experiments were purchased from Janvier Labs (Le Genest-Saint-Isle). For tissue distribution and measurements studies BALB/cJ (8-9 weeks old) mice weighing 23-31 g at the time of experiments were ordered from Jackson Laboratories (Bar Harbor, Maine USA). The BALB/cJ strain of mice was chosen for its highly anxious phenotype (Dulawa et al., 2004) and was used for plasma and brain tissue levels of ketamine and its metabolites, for behavioral and microdialysis studies performed 24h post-dose. Male BALB/cJ and CD-1 mice (9-10 weeks old) weighing 30-50 g at the beginning of the experiments were purchased from Charles River Laboratories (Raleigh, NC, USA) and were used for the pharmacokinetic (PK) study from 0 to 4h post-dose because of prior studies revealing their sensitivity to behavioral actions of ketamine and hydroxynorketamines (Zanos et al., 2016). Protocols were approved by the Institutional Animal Care and Use Committee in France (Council directive # 87-848, October 19, 1987, “*Ministère de l’Agriculture et de la Forêt, Service Vétérinaire de la Santé et de la Protection Animale, protocol APAFIS #5489, permissions # 92-196*” to A.M.G.) as well as with the European directive 2010/63/EU, or the University of Maryland, Baltimore Animal Care and Use Committee and were conducted in full accordance with the National Institutes of Health Guide for the Care and Use of Laboratory Animals.

All mice were housed 4-5 per cage in a room with a controlled temperature (21 ± 1°C) on a 12-h/12-h light/dark cycle and were tested during the light phase. Food and water were available *ad libitum* except during behavioral observations. For microdialysis experiments, mice were single-housed after the probes were placed. Efforts were made to minimize the number of mice used in the experiments.

### 2.2. Drugs

(*R*,*S*)-ketamine and fluconazole were purchased from Sigma-Aldrich (Saint-Quentin Fallavier, France, and St. Louis Missouri, USA). (*2R,6R*)-hydroxynorketamine [(*2R,6R*)-HNK] was provided by the National Center for Advancing Translational Sciences (Bethesda, MD, USA). The CYP inhibitor fluconazole (Fresenius Kabi; 10 mg/kg and 20 mg/kg) was diluted in vehicle (NaCl 0.9%) and administered intraperitoneally (i.p.) 1 h before ketamine administration (10 mg/kg, i.p.) (Uchida et al., 2014; Yang et al., 1996); *in vivo* microdialysis and behavioral tests were performed 24h after ketamine administration. Fluconazole is a potent inhibitor of CYP2C9 and a moderate inhibitor of CYP3A4 and CYP2C19 in humans and rodents (FDA, 2019; Gupta et al., 2007; Uchida et al., 2014). Drug doses and pre-treatment times were based on previous studies performed with ketamine (Pham et al., 2018; Pham et al., 2017).

### 2.3. Experimental design

Experimental timelines are shown in Fig. 1a, 2a, 3a, 4a and 5a. For the pharmacokinetic studies, plasma and brains were collected at 2.5, 5, 10, 30, 120, and 240 min after ketamine administration according to published methods (Zanos et al., 2016). For the assessment of ketamine and metabolite levels at t24h in BALB/cJ mice, plasma samples were collected from the sub-mandibular vein immediately after behavioral testing, as previously described (Mendez-David et al., 2013). Behavioral testing occurred during the light phase between 08:00 and 12:00. Then, animals were sacrificed and the prefrontal cortex was dissected to quantify the levels of (*R*,*S*)-ketamine and its metabolites.

**Fig. 1.**
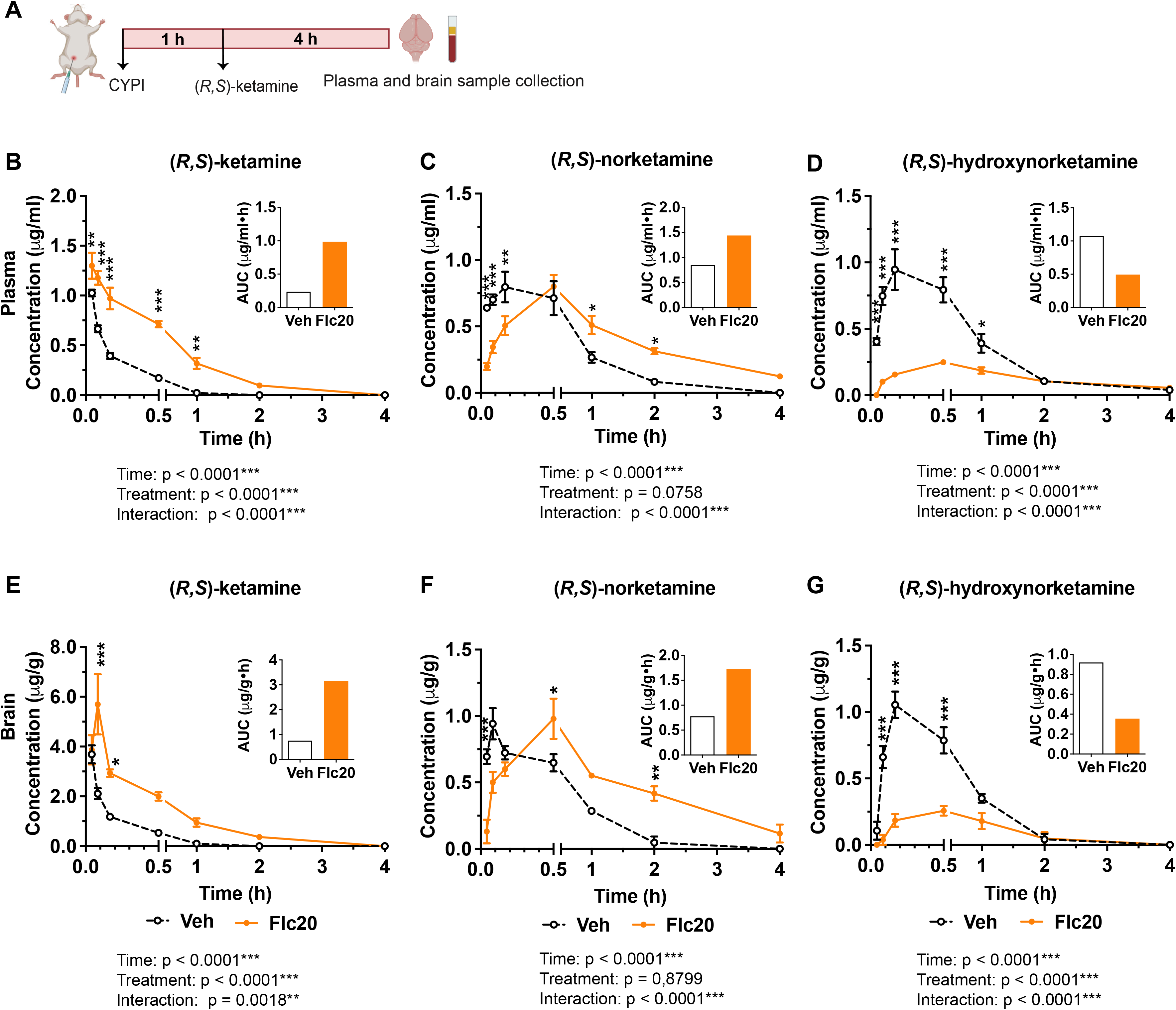
CYPI pre-treatment increases plasma and brain levels of (R,S)-ketamine and (R,S)-norketamine and decreases those of (R,S)-HNK within the first 4 hours post-treatment. (A) Experimental design timeline. (*R,S*)-ketamine (10 mg/kg, i.p.) was administered 1 hour after fluconazole administration (20 mg/kg, i.p.). Plasma and brain samples were collected for 4 hours. Pharmacokinetic profiles of (*R,S*)-ketamine, (*R,S*)-norketamine and (*R,S*)-HNKs in BALB/cJ mice (B-G)). Mean ± S.E.M. (number of mice /group = 10). *p<0.05, **p<0.01 and ***p<0.001 versus Veh/(*R,S*)-ket. CYPI: cytochrome P450 enzyme inhibitor; Veh: vehicle; Flc: fluconazole.

**Fig. 2.**
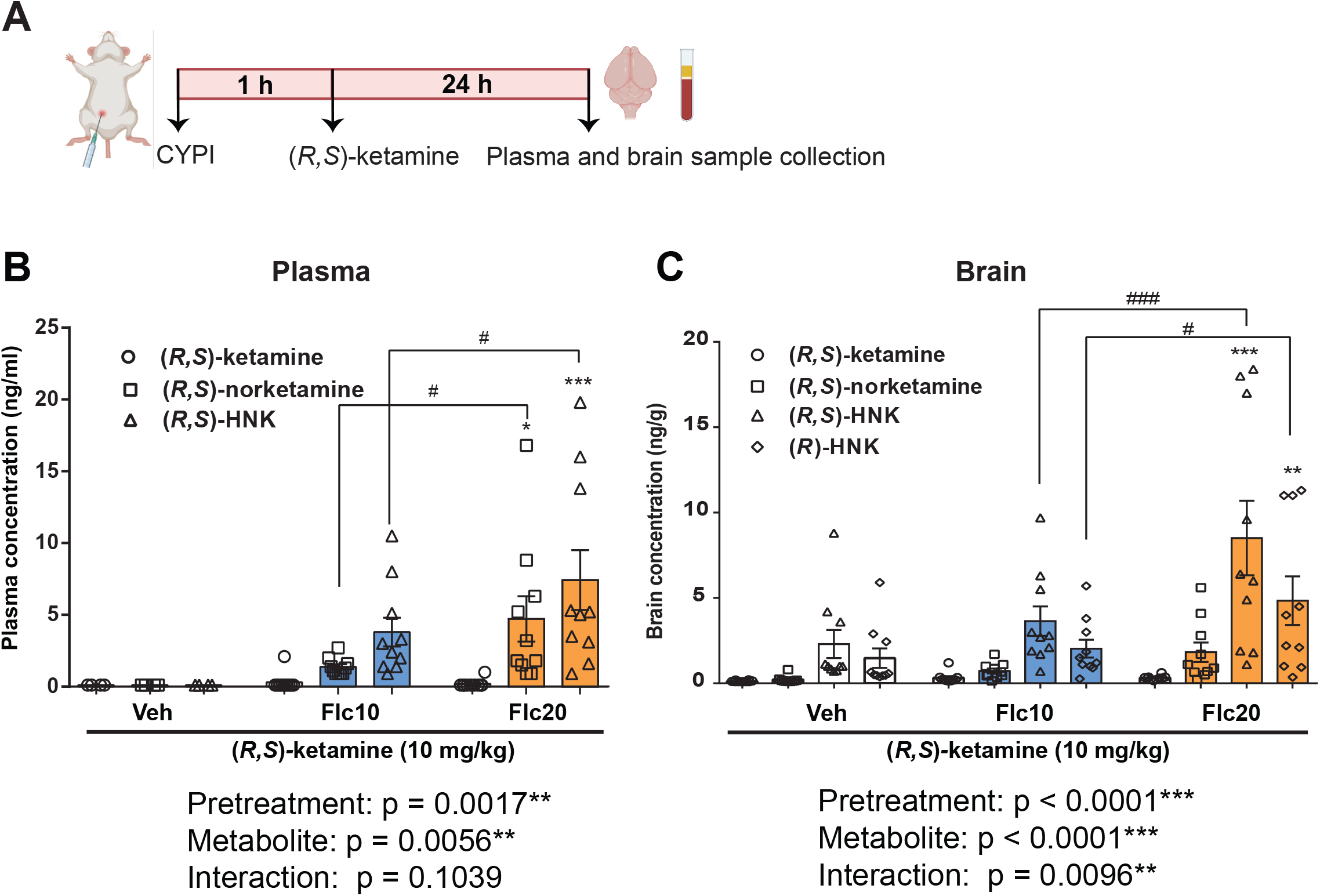
Trace concentrations of (R,S)-norketamine and (6)-HNK were measured in the plasma and brain 24h after CYPI pre-treatment. (A) Experimental design timeline. *(R,S)*-ketamine (10 mg/kg, i.p.) was administered 1 hour after fluconazole administration (10 and 20 mg/kg i.p.). Plasma samples were collected at t24h from the sub-mandibular vein of each individual mouse. Ketamine and its metabolite levels in plasma (B) and brain (C) at t24h. Fluconazole alone had no significant effect on the plasma and brain *(R,S)*-ketamine levels at t24h post-ketamine administration. Mean ± S.E.M. (number of mice/group = 10). *p<0.05, **p<0.01 and ***p<0.001 *versus* Veh/Ket. #p<0.05, ###p<0,001 *versus* the Flc10/Ket. CYPI: cytochrome P450 enzyme inhibitor; Veh: vehicle; Flc: fluconazole; Ket: ketamine

**Fig. 3.**
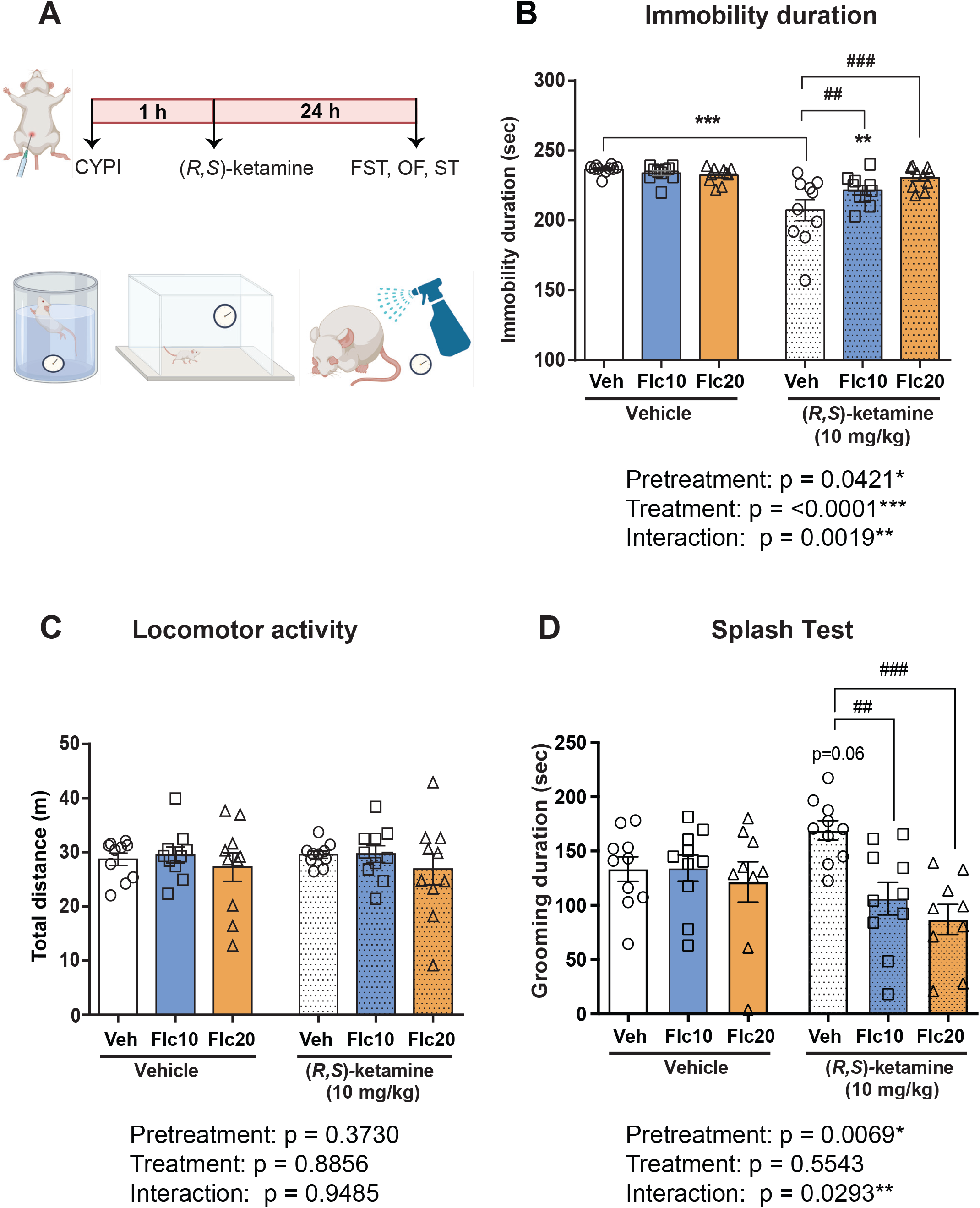
CYPI pre-treatment blocks the sustained antidepressant-like activity of (R,S)-ketamine in BALB/cJ mice. Fluconazole (Flc; 10 and 20 mg/kg i.p.) was administered 1 h before *(R,S)*-ketamine (10 mg/kg i.p.): (A) Experimental design for the studies following the systemic administration of *(R,S)*-ketamine (10 mg/kg, i.p.); (B) immobility duration in the forced swim test (FST); (C) locomotor activity in the open field test (OF). (D) grooming duration in the splash test. Fluconazole prevented the sustained antidepressant-like activity of *(R,S)*-ketamine at t24h in the FST and the splash test. The locomotor activity was not altered in any of the groups. Mean ± S.E.M. (number of mice/group = 10). **p<0.01 and ***p<0.001 *versus* Veh/Veh controls. ##p<0.01 and ###p<0.001 *versus* the Veh/*(R,S)*-Ket. CYPI: cytochrome P450 enzyme inhibitor; Veh: vehicle; Flc: fluconazole; Ket: ketamine.

**Fig. 4.**
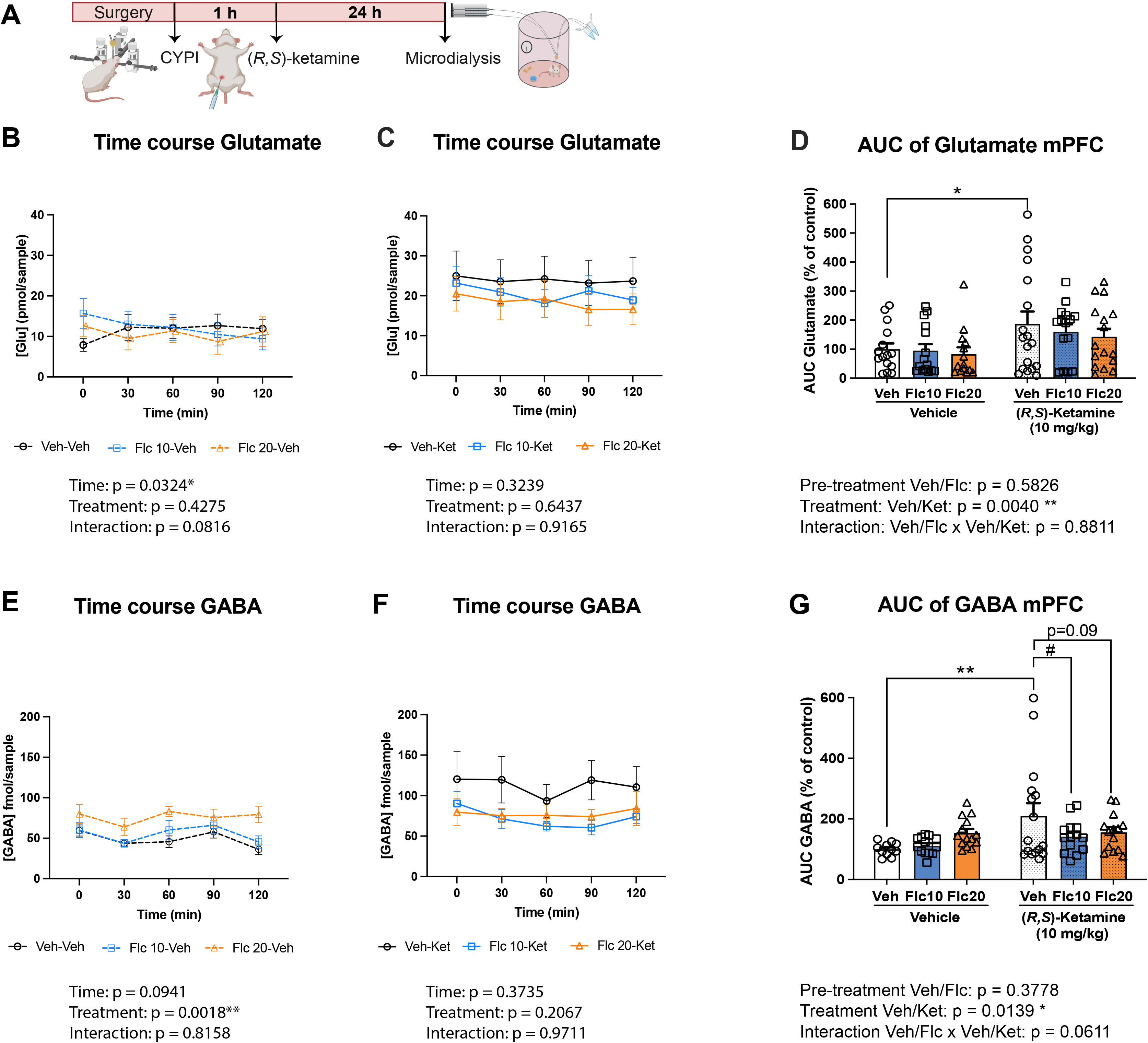
CYPI pre-treatment decreases (R,S)-ketamine-induced glutamate and GABA release in the mPFC in BALB/cJ mice. (A) Experimental design timeline. (B, C, E, F) Time course of changes in extracellular glutamate and GABA levels from t0-t120 min as measured 24 hours after ketamine administration. (D, G) AUC values of extracellular glutamate (Glu) and GABA levels in the mPFC. Dialysate samples were collected from 0 to 120 min and presented in pmol/sample (Glu) and fmol/sample (GABA). AUC values of Glu and GABA were calculated by the linear trapezoidal method and presented in percentages of the Veh/Veh group. Fluconazole, (10 i.p. given 1 h before ketamine), decreased ketamine-induced increases in [GABA]ext in the mPFC at t24h post-ketamine administration. Mean ± S.E.M. (number of mice/group = 10). *p<0.05, **p<0.01 vs Veh/Veh. #p<0.05 vs Veh/Ket. CYPI: cytochrome P450 inhibitor; Veh: vehicle; Flc: fluconazole; Ket: ketamine.

### 2.4. Behavioral assessment

The antidepressant-like and exploratory/anxiolytic responses elicited by ketamine administered 24 h before behavioral testing were assessed in the Forced Swim Test (FST), Open Field Test (OF), or Splash Test. Separate cohorts of mice were tested in each behavioral paradigm.

#### 2.4.1. Forced Swim Test (FST)

The FST procedure was modified to enhance the sensitivity for detecting the putative antidepressant-like activity of drugs (Holick et al., 2008; Porsolt et al., 1977). Mice were placed into clear plastic cylinders (15 cm in diameter and 23 cm deep), filled two-thirds with water at 23–25◦C for 6 min. Immobility time was measured during the last 4 min of the test.

#### 2.4.2. Open Field (OF) test

Motor activity in the OF was quantified in 39 × 39 cm perpex plastic open field boxes (Vivo-tech/Ugo Basile, Salon de Provence, France). The apparatus was illuminated from the ground with specially designed 40 x 40 cm Infra-red backlights (monochromatic wavelength 850 nm high homogeneity, Vivo-tech, Salon de Provence, France). Testing was performed over a period of 10 min, during which the locomotor activity of the mouse was recorded using infrared cameras (Vivo-tech, Salon de Provence, France). Total ambulatory distance over 10 min after systemic administration was analyzed using ANYMAZE version 6 video tracking software (Stoelting Co/Vivo-tech, Salon de Provence, France) (Faye et al., 2019).

#### 2.4.3. Splash test

A 10% sucrose solution was applied to the mouse’s snout, and the duration and latency of grooming were manually recorded, as previously described (David et al., 2009; Mendez-David et al., 2014).

### 2.5. Intracerebral in vivo microdialysis in the medial prefrontal cortex

Each mouse was anesthetized using chloral hydrate (400 mg/kg, i.p.) and implanted bilaterally with 2 microdialysis probes (CMA7 model, Carnegie Medicine, Stockholm, Sweden) located in the medial prefrontal cortex (mPFC). Stereotaxic coordinates were as follows (mm from bregma): mPFC: A= + 2.2, L= ± 0.5, V= - 3.4; (A, anterior; L, lateral; and V, ventral) (Nguyen et al., 2013; Pham et al., 2018; Pham et al., 2017). On the same day, after awakening, mice received an acute injection of vehicle or fluconazole (10 mg/kg or 20 mg/kg, i.p.). One hour later, the animals received an i.p. injection of saline or (*R*,*S*)-ketamine (10 mg/kg). The next day, ∼24h after ketamine administration, the probes were continuously perfused with an artificial cerebrospinal fluid (aCSF, composition in mmol/L: NaCl 147, KCl 3.5, CaCl_2_ 1.26, NaH_2_PO_4_ 1.0, pH 7.4 ± 0.2) at a flow rate of 1.0 µl/min through the mPFC using CMA/100 pump (Carnegie Medicine, Stockholm, Sweden), while mice were awake and freely moving in their cage. One hour after the start of aCSF perfusion stabilization period, four fractions were collected (one every 30 min) to measure the extracellular levels of glutamate, GABA and glutamine ([Glu]_ext_ and [GABA]_ext_ in the mPFC by liquid chromatography-tandem mass spectrometry (LC-MS/MS) (lower limit of quantification LLOQ = 2 ng/ml: (Defaix et al., 2018). AUC values (% of baseline) for the 0-120 min period of samples’ collection were also calculated as previously described (Nguyen et al., 2013). At the end of the experiments, localization of microdialysis probes was verified histologically (Bert et al., 2004). Animals were removed if probes were not in the correct mPFC location.

### 2.6. Measurement of plasma and brain concentrations of (R,S)-ketamine, (R,S)-norketamine and (6)-HNK

Male CD-1 (PK study from 0 to 4 h post-dose) and BALB/cJ (PK study from 0 to 4 h post-dose and at 24h post-dose) mice received a fluconazole (20 mg/kg; i.p.) or vehicle (saline; i.p.) injection followed one hour later by ketamine (10 mg/kg; i.p.) injection. Mice were deeply anesthetized with 3-4% isoflurane (2 min exposure) and decapitated at 2.5, 5, 10, 30, 60 or 240 min post-ketamine injection. Trunk blood was collected in EDTA-containing tubes and centrifuged at 8000 x g for 6 min at 4°C. Plasma was collected and stored at −80°C until analysis. Whole brains were simultaneously collected, immediately frozen on dry ice, and stored at −80°C until analysis.

Achiral liquid chromatography–tandem mass spectrometry was used to determine the concentration of ketamine and its metabolites in plasma as previously described (Farmer et al., 2020) and in brain with slight modifications (Farmer et al., 2020). Briefly, brains were suspended in 480 μl of water:methanol (3:2, v/v) with the addition of d_4_-ketamine (10 μl of 10 μg/ml) and d_4_-norketamine (10 μl of 10 μg/ml). The subsequent mixture was homogenized on ice and centrifuged at 21000 × *g* for 30 min. The supernatant was processed using 1-ml Oasis HLB solid-phase extraction cartridges (Waters Corp., Waltham, MA) and transferred to an autosampler vial for analysis. Quality control standards were prepared at 156.25 ng/ml, 312.5 ng/ml and 1250 ng/ml for plasma and 250 ng/ml for brain. The calibration standards for (*R,S*)-ketamine, (*R,S*)-norketamine and (*2R,6R*;*2S,6S*)-HNK ranged from 19.53 to 5,000 ng/ml for both plasma and brain. MS/MS analysis was performed using a triple quadropole mass spectrometer model API 4000 sytstem (Applied Biosystesms/MDS Sciex) equipped with Turbo Ion Spray (TIS). The quantification of (*R,S*)-ketamine, (*R,S*)-norketamine, and HNK levels was accomplished by calculating area ratios using d_4_-ketamine as the internal standard for plasma and ketamine/d_4_-ketamine for ketamine *(R,S)*-norketamine and HNK/d_4_-norketamine for norketamine and HNK was calculated using Graphpad Prism v8.21.

A liquid chromatography–tandem Endura triple quadrupole mass spectrometer (Thermo Fischer^®^, Les Ulis, France) was used with ketamine-d4 as internal standard to determine the levels of (*R*,*S*)-ketamine, (*R,S*)-norketamine, and (*2R,6R*;*2S,6S*)-HNK at t24h post-ketamine injection in the plasma and the brain tissues as previously described (Abe et al., 2013; Larabi et al., 2023; Pham et al., 2018). The method was fully validated for selectivity, carry-over, linearity, limit of detection (LOD), limit of quantification (LOQ), accuracy, precision, matrix effect, and extraction recovery. The limits of detection (LOD) and quantification (LOQ) were 0.1 ng/ml and 0.1 ng/g for the plasma and brain, respectively (Abe et al., 2013; Larabi et al., 2023).

### 2.7. Statistical Analyses

All experimental results are given as the mean ± S.E.M. Data were analyzed using Prism *v*10.0.2 software (GraphPad, San Diego, CA, USA). Data analyses and comparisons between groups were performed using either an unpaired t-test or a one-way ANOVA or a two-way ANOVA followed by an appropriate *post-hoc* analysis. Since there is an a priori rationale to assess whether inhibition of (*R,S*)-ketamine’s metabolism affects ketamine-induced neurotransmitter changes, thus affecting antidepressant-relevant behavioral actions, we performed planned comparison tests wherever pre-treatment and treatment factorial ANOVAs were significant. Tissue distribution data for up to 4 hours post-treatment were analyzed using two-way ANOVA with factors time and pre-treatment (vehicle *versus* fluconazole) followed by Fisher’s LSD*post-hoc* analysis. Statistical significance was set at *p* ≤ 0.05. A summary of statistical measures is included in Supplementary *Table S1-Statistics*.

## 3. Results

### 3.1. CYPI pre-treatment increases plasma and brain exposure to (R,S)-ketamine and (R,S)-norketamine and decreases those of HNK within the first 4 hours post-treatment

The plasma and brain levels of ketamine, (*R,S*)-norketamine, and (*2R,6R;2S,6S*)-HNK were monitored during the first 4 hours after ketamine treatment (10 mg/kg, i.p.) in BALB/cJ and CD-1 mice pre-treated with vehicle or fluconazole (20 mg/kg, i.p.). In both strains, pre-treatment with fluconazole robustly attenuated metabolism of ketamine to (*2R,6R;2S,6S*)-HNK, resulting in greater total exposure to ketamine and (*R,S*)-norketamine, and lower total exposure to (*2R,6R;2S,6S*)-HNK in the plasma and the brain, relative to vehicle pre-treatment (Fig. 1; inset, AUC). In the fluconazole-treated mice, plasma concentrations of ketamine were higher during the first 1 hour in BALB/cJ mice (Fig. 1B) and during the first 30 minutes after ketamine administration in CD-1 (Fig. S1B), relative to vehicle-treated controls. Plasma norketamine levels were initially lower in fluconazole-treated mice (i.e. 2.5-10 minutes after ketamine dosing), similar at 30 minutes, and higher at later time points (i.e. 1-2 h in BALB/cJ mice and 1 h in CD-1 mice (Fig. 1C and S1C), compared to the vehicle group. In BALB/cJ mice, plasma ketamine and norketamine remained above LOQ for longer in fluconazole-treated mice compared with controls (Fig. 1B, 1C). Finally, fluconazole pre-treatment resulted in lower plasma concentrations of (*2R,6R;2S,6S*)-HNK over the first hour after ketamine administration in both strains (in BALB/cJ mice, Fig. 1D; in CD-1 mice, Fig. S1D). In both strains, fluconazole treatment did not alter the time at which peak plasma concentrations were observed (Tmax) for (*R,S*)-ketmaine (2.5 min), but it delayed the Tmax for (*R,S*)-norketamine and (*2R,6R;2S,6S*)-HNK (10 min in vehicle-treated mice *versus* 30 min in fluconazole-treated mice) (Fig. 1). Thus, fluconazole both delayed the formation of (*2R,6R;2S,6S*)-HNK concentrations and attenuated its peak plasma concentrations and total exposure.

In the brain, total exposure to ketamine and its initial concentrations (i.e., between 5-10 min in BALB/cJ, Fig. 1E, mice and ≤ 10 min in CD-1 mice, Fig. S1E) were higher in fluconazole-treated mice. Total brain exposure to (*R,S*)-norketamine and its initial concentrations (≤ 5 min in BALB/cJ mice, Fig. 1F, and ≤ 10 min in CD-1 mice, Fig. S1F) were lower in fluconazole-treated mice, but at later time points (at 30 min and 2 h in BALB/cJ mice, and at 30 min-1 h in CD-1 mice) fluconazole treatment resulted in higher (*R,S*)-norketamine levels, compared to vehicle-treated controls. Total exposure to and initial concentrations (i.e., ≤ 30 min in BALB/cJ mice, Fig. 1G and ≤ 1 hr in CD-1 mice, Fig. S1G) of (*2R,6R;2S,6S*)-HNK in the brain were attenuated in fluconazole-treated mice. In the brain, fluconazole treatment resulted in a later Tmax observed for (*R,S*)-ketamine (5 min for fluconazole vs. 2.5 min for vehicle), (*R,S*)-norketamine (30 min for fluconazole vs. 5 min for vehicle), and (*2R,6R;2S,6S*)-HNK (30 min for fluconazole vs. 10 min for vehicle) in both strains (Fig. 1). In both the plasma and the brain, levels of additional HNK metabolites were below the limits of quantification in all groups between 2.5 min and 4 hours post-ketamine dosing.

### 3.2. Trace concentrations of (R,S)-norketamine and (6)-HNKs were detected in the plasma and brain 24 h after fluconazole/ketamine

In the vehicle pre-treated group, plasma concentrations of (*R*,*S*)-ketamine and (*2R,6R;2S,6S*)-HNKs were close to the limit of detection (LOD) at t24h, consistent with our previous findings (Pham et al., 2018) (Fig. 2B). However, in fluconazole pre-treated groups, plasma concentrations of (*R*,*S*)-norketamine and (*6*)-HNKs were quantifiable at t24h (Fig. 2B). The fluconazole-induced increase in (*R,S*)-norketamine and (*6*)-HNKs showed a dose-response, with the highest dose of fluconazole (20 mg/kg) resulting in significantly greater plasma levels of (*R,S*)-norketamine and (*6*)-HNKs compared to the lower (10 mg/kg) dose.

A two-way ANOVA shows a statistically significant difference in plasma concentrations of (*R,S*)-norketamine (*p*<0.05) and (*6*)-HNKs (*p*<0.001) at t24h in mice pre-treated with fluconazole 20 mg/kg compared to those of the ketamine-treated group (Fig. 2B). However, these (*6*)-HNKs levels correspond to approximately 0.5% of the plasma concentration previously found at 30 min post dose in BALB/cJ mice (see here in Fig. 1D and in Fig. 1E in (Pham et al., 2018)). In addition, a statistically significant increase in plasma concentrations of (*6*)-HNKs (*p*<0.05) was found at t24h in mice pre-treated with fluconazole 20 mg/kg compared to those pre-treated with fluconazole 10 mg/kg (Fig. 2B).

For the brain samples, a chiral separation column was used to more precisely quantify the concentrations of HNKs. In the vehicle pre-treated group, brain concentrations of (*R*,*S*)-ketamine, (*R,S*)-norketamine, and (6)-HNKs were close to the LOD at t24h. However, in fluconazole-pre-treated mice, brain concentrations of (*6*)-HNKs and (*R*)-HNK at t24h were still quantifiable (Fig. 2C). A two-way ANOVA shows a statistically significant difference [F(3,120)=11.59; p<0.001] in brain concentrations of (*6*)-HNKs (p<0.001) and (*R*)-HNK (p<0.01), compared to ketamine-treated group (Fig. 2C). In addition, brain concentrations of (*6*)-HNKs (*p*<0.001) and (*R*)-HNK (*p*=0.05) at t24h in mice pre-treated with fluconazole 20 mg/kg were higher to those pre-treated with fluconazole 10 mg/kg (Fig. 2C). No differences were found for the concentrations of (*R,S)*-norketamine (Fig. 2C).

### 3.3. CYPI pre-treatment blocks the sustained antidepressant-like behavioral activity of (R,S)-ketamine in mice 24h post-dose

We tested if pre-treatment with fluconazole alters ketamine’s behavioral effects 24 hours after administration (Fig. 3A). Analysis of the immobility duration in the FST using a two-way ANOVA revealed a significant effect of fluconazole pre-treatment [F(2,54)=3.362, *p*=0.042], a significant effect of ketamine [F(1,54)=23.20, *p*<0.0001] and a significant interaction between these two factors [F(2,54)=7.055, *p*=0.0019] (Fig. 3B). Thus, a single dose of ketamine decreased the immobility duration compared to vehicle group (p<0.001). Fluconazole (10 or 20 mg/kg, i.p.) 1h before ketamine administration, prevented the effect of ketamine on immobility duration (p<0.001) (Fig. 3B).

Analysis of the locomotor activity in the OF test using a two-way ANOVA revealed no statistically significant differences in the total distance traveled among the six groups studied regarding the pre-treatment factor [F(2,54)=1.004; *p*=0.373], the treatment factor [F(1,54)=0.021; *p*=0.8856] and no interaction between these factors [F(2,54)=0.05; *p*=0.9485] (Fig. 3C).

Analysis of the grooming duration in the splash test using a two-way ANOVA revealed a significant effect of fluconazole pre-treatment [F(2,51)=5.494; *p*=0.0069], no significant effect of ketamine [F(1,51)=0.3544, p=0.5543] and a significant interaction between these factors [F(2,51)=3.784, *p*=0.0293] (Fig. 3D). Thus, fluconazole (10 or 20 mg/kg, i.p.) 1h before ketamine administration, prevented ketamine’s effect on grooming duration (p=0.0013 and p=0.0004, respectively) (Fig. 3D).

Similarly, in CD-1 mice, analysis of the immobility duration in the FST using a two-way ANOVA revealed a significant effect of fluconazole pre-treatment [F(1,36)=9.467, *p*=0.0040], a significant effect of ketamine [F(1,36)=15.20, *p*<0.001] and a significant interaction between these two factors [F(1,36)=9.493, *p*=0.0039] (Fig. S1H). While ketamine (10 mg/kg, i.p.) reduced immobility duration in mice pre-treated with saline (p<0.0001, supplemental Fig. S1H), pre-treatment with fluconazole (20 mg/kg, i.p.) blocked this effect (p=0.0001, supplemental Fig. S1H). Prior treatment with fluconazole did not alter immobility time within the vehicle group (Supplemental Fig. S1H).

### 3.4. CYPI pre-treatment decreases ketamine-induced neurotransmitter release in the mPFC 24h post-dose

Finally, given the well-established link between increased glutamate release in the mPFC and antidepressant-like activity of ketamine, we performed *in vivo* microdialysis experiments to test if pre-treatment with fluconazole alters ketamine’s neurochemical effects in the mPFC, 24 hours after administration in BALB/cJ mice (protocol in Fig. 4A). The average neurotransmitter basal values B0 at t0 in the mPFC for six groups are summarized in Table S2.

Cortical extracellular levels of glutamate [Glu]_ext_ and GABA [GABA]_ext_ in the Flc10/Veh and Flc20/Veh groups were not statistically different from the Veh/Veh group (Fig. 4D, 4G).

Changes in extracellular glutamate levels were analyzed in Fig. 4B, 4C, 4D). A two-way ANOVA on AUC values (% of controls) revealed no significant effect of fluconazole pre-treatment [F(2,87)=0.544; *p*=0.5826], a significant effect of ketamine [F(1,87)=8.719, p=0.0040] and no significant interaction between these factors [F(2,87)=1.127, *p*=0.8811] (Fig. 4D). Planned comparison tests indicated that ketamine significantly increased AUC values of [Glu]_ext_ as compared to vehicle-treated mice (Fig. 4D). However, cortical [Glu]_ext_ in the fluconazole/ketamine group did not differ from vehicle-treated mice at t24h post-treatment (Fig. 4D).

Changes in extracellular GABA levels were analyzed in Fig. 4E, 4F, 4G). A two-way ANOVA on AUC values (% of controls) revealed no significant effect of fluconazole pre-treatment [F(2,76)=0.986; *p*=0.3778], a significant effect of ketamine [F(1,76)=6.347, p=0.0139] and a trend for an interaction between these factors [F(2,76)=2.90, *p*=0.061] (Fig. 4G). Planned comparison tests indicated that ketamine significantly increased AUC values of [GABA]_ext_ as compared to vehicle-treated mice. In addition, [GABA]_ext_ in the Flc10/ketamine-treated group was reduced as compared to ketamine-treated mice, while a trend to [GABA]_ext_ decrease was measured in the Flc20/ketamine group as compared to ketamine-treated mice at t24h post-treatment (Fig. 4G).

### 3.5. (2R,6R)-HNK rescues the sustained antidepressant-like behavioral activity of (R,S)-ketamine in CYPI-induced blockade 24h post-dose in BALB/cJ mice

We tested if (*2R,6R*)-HNK administration can reverse fluconazole-induced blockade of ketamine’s behavioral effects 24 hours after administration (experimental protocol, Fig. 5A). A one-way ANOVA revealed statistically significant differences between treated-groups in the FST [F(3,34)=4.13, *p*=0.0134]. Ketamine decreased immobility duration compared to the vehicle-treated mice (\**p*<0.05) as already described in Fig. 3B. Fluconazole (20 mg/kg, i.p.) 1h before ketamine administration (10 mg/kg, i.p.), prevented the effect of ketamine on immobility duration (##p<0.01) (Fig. 5B). Finally, a single administration of (*2R,6R*)-HNK rescued the antidepressant-like effect of ketamine in mice pretreated with fluconazole ($p<0.05) (Fig. 5B).

**Fig. 5.**
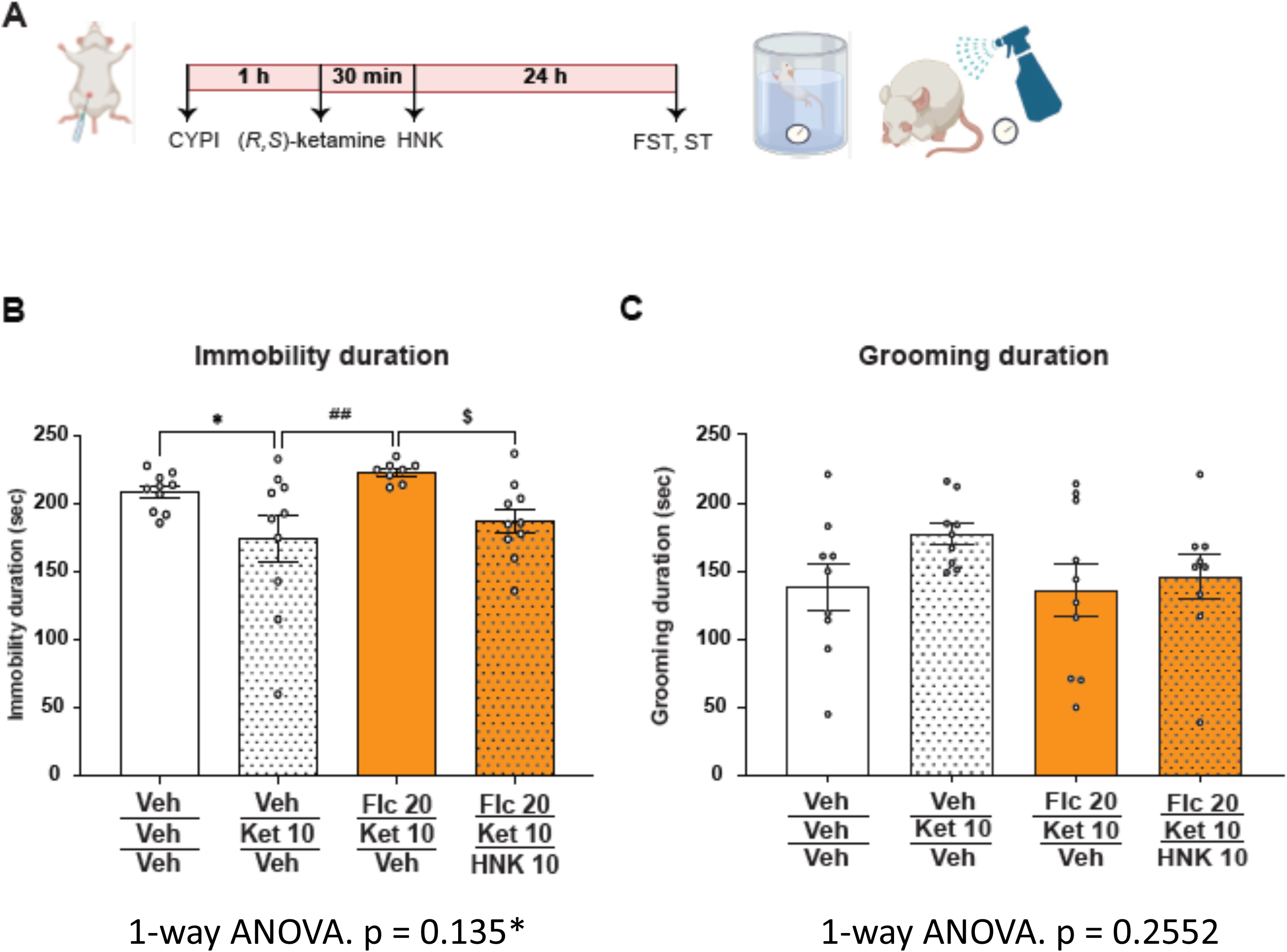
(2R,6R)-HNK rescues the sustained antidepressant-like activity of (R,S)-ketamine in CYPI-treated mice 24h post-dose in BALB/cJ mice. Fluconazole (Flc; 20 mg/kg i.p.) was administered 1 h before (*R,S*)-ketamine (10 mg/kg i.p.) and (*2R,6R*)-HNK (10 mg/kg i.p.) was administered 30 minutes after (*R,S*)-ketamine (10 mg/kg i.p.). (A) Experimental design for the studies following the systemic administration of the three molecules; (B) (*R,S*)-Ketamine decreased immobility duration, and fluconazole prevented its sustained antidepressant-like activity at t24h in the forced swim test (FST), *p<0.05 Veh/ket10 *versus* the Veh/control group. A fluconazole pre-treatment blocked ketamine’s antidepressant-like effects in the FST, ##p<0.01 Veh/Ket10 *versus* Flc20/ket10. A single administration of (*2R,6R*)-HNK rescued the antidepressant-like activity of ketamine in mice pretreated with fluconazole, $p<0.05 Flc20/Ket10/HNK10 *versus* Flc20/Ket10. Mean ± S.E.M. (number of mice/group = 10). CYPI: cytochrome P450 enzyme inhibitor; Veh: vehicle; Flc: fluconazole; Ket: ketamine; HNK: (*2R,6R*)-HNK. (C) The grooming duration in the sucrose ST did not reveal statistically significant differences between treated groups.

Analysis of the immobility duration in the sucrose ST using a one-way ANOVA did not show statistically significant differences between treated-groups [F(3,33)=1.417, *p*=0.2552] (Fig. 5C).

## 4. Discussion

(*2R,6R*)-HNK, one of the major brain metabolites of (*R,S*)-ketamine has been proposed to be essential for the sustained antidepressant-like activity of ketamine (Suzuki et al., 2017; Zanos et al., 2016). Conversely, it was claimed that the metabolism to (*2R,6R; 2S,6S*)-HNK [(*6*)-HNK)] is not necessary to induce this effect (Yamaguchi et al., 2018). Here, we sought to clarify these discrepancies by assessing the behavioral and neurochemical consequences of inhibiting hepatic cytochrome P450 (CYP) enzymes, thereby attenuating (*R*,*S*)-ketamine metabolism to (*6*)-HNK. Surprisingly, a combined CYP inhibitor approach with ketamine has been little investigated in rodents to date.

We characterized the specific role of ketamine’s metabolism in its antidepressant actions. We evaluated the behavioral and neural effects of a pre-treatment with fluconazole, a potent inhibitor of hepatic cytochrome P450 enzymes on antidepressant responses of ketamine. We discovered that fluconazole altered the pharmacokinetic profile of ketamine, by increasing both plasma and brain levels of ketamine and (*R*,*S*)-norketamine, while robustly reducing those of (*6*)-HNKs. Furthermore, fluconazole pretreatment attenuated the sustained antidepressant-like behavioral actions of ketamine that appeared at the same time (t24h) as the attenuation in ketamine-induced neuronal release of GABA in the mPFC. Overall, our results are consistent with an essential role of (*6*)-HNKs in mediating those sustained actions of ketamine. To our knowledge, potential interactions between CYPIs and ketamine during antidepressant treatment has not previously been studied in major depressive disorder (MDD) patients.

Here, our tissue distribution and clearance data measured following administration of a single dose of ketamine demonstrate that pre-treatment with fluconazole, a CYPI administered prior to ketamine, robustly attenuated the metabolism of ketamine to (6)-HNK, concomitantly resulting in greater plasma and brain exposure to (*R*,*S*)-ketamine and (*R*,*S*)-norketamine and lower concentrations of (6)-HNK during the 4 hour period after (*R,S*)-ketamine dosing. Concentrations of (*R,S*)-norketamine in fluconazole-treated mice are initially lower in the first 30 min to 1 h, then higher at later time points, consistent with an early attenuation of both the metabolism of (*R,S*)-ketamine to (*R,S*)-norketamine and that of (*R,S*)-norketamine to (*6*)-HNK. Additionally, our experiments reveal that fluconazole pretreatment results in very low, but measurable levels of (*R,S*)-norketamine and (*R,S*)-HNK remaining in the plasma and brain for an extended period of time including at t24h, further indicating that metabolism to (*6*)-HNK was robustly delayed. Interaction of a CYPI with a drug is expected to increase the levels of the parent drug (Can et al., 2016; Palkama et al., 1998), while decreasing those of its metabolites, consistent with our findings here: ketamine lost its antidepressant-like effects in mice pre-treated with a CYP inhibitor suggesting that (*6*)-HNK metabolites contribute to the sustained antidepressant-like activity of ketamine, at least under our experimental conditions.

One study investigated the role of the metabolism of the isomers, (*R*)-ketamine, not the racemic mixture, in its antidepresssant-like activity (Yamaguchi et al., 2018). Pharmacokinetic profile and acute behavioral effects of (*R*)-ketamine in the presence of CYP inhibitors suggests that (*2R,6R*)-HNK is not essential for the antidepressant actions of (R)-ketamine in mice (Yamaguchi et al., 2018): these results contradict those of the present study. Comparison of experimental conditions between our study and Yamaguchi et al., (2018) (Yamaguchi et al., 2018) (Table 1) show differences in stress-induced phenotype of mice (LPS-induced inflammation model or in C57BL/6 naive mice), pre-treatment with a cocktail of CYP inhibitors (ticlopidine and 1-ABT), a high (*R*)-ketamine dose (30 mg/kg) and short time points (but not at t24h) for the evaluation of (*R*)-ketamine effects. However, CYPI prolonged the plasma elimination half-life t_1/2_ of (*R*)-ketamine and (*R*)-norketamine up to 8 hours post-treatment (Yamaguchi et al., 2018) and increased C_max_ and AUC_0-3hr_, which is consistent with our findings. Among these experimental conditions, differences in the main time point of observation (at 3hr *versus* t24h post-dose) may explain these discrepancies between Yamaguchi et al., (2018) and the present results. By contrast, when (*R*)-ketamine was compared with a deuterated form of the drug, which hinders its metabolism to (*2R,6R*)-HNK (Zanos et al., 2019), it was found that metabolism of (*R*)-ketamine to (*2R,6R*)-HNK increased the potency of (*R*)-ketamine to exert antidepressant-relevant actions in mice.

To our knowledge the two blunted effects of ketamine on antidepressant-like activity and mPFC [GABA]_ext_ at t24h following a CYPI pre-treatment in a BALB/cJ model of anxiety/depression was observed here, for the first time. In addition, the results of the rescue experiment are also new. The antidepressant-like effect of ketamine was absent in fluconazole-pre-treated mice 24 hours after treatment. By contrast, a single administration of the metabolite (*2R,6R*)-HNK was sufficient to rescue ketamine antidepressant-like activity in the FST 24 hours following treatment in the fluconazole-treated group. In addition, our behavioral data in Fig. 3 has no evidence that fluconazole is having non-specific effects, i.e., no effects by itself in any of our behavioral models.

Overall, the present results rule out a non-specific action of the CYPI fluconazole, and confirm that metabolism from ketamine to (*6*)-HNKs by CYP enzymes is necessary for the full antidepressant-like effects of the parent drug when administered to rodents. They contradict those of Yamaguchi et al., (2018) showing that metabolism to (*2R,6R*)-HNK is not essential for the antidepressant effects of (*R*)-ketamine, one of the enantiomers of (*R,S*)-ketamine in mice (Yamaguchi et al., 2018).

For rapid-acting antidepressant, both an induction/initiation, as well as expression mechanisms of antidepressant action have been described, and are distinct phenomena (Gould et al., 2019). The induction mechanism occurs immediately after drugs’ exposure, while the expression mechanism is observed at a time point, e.g., t24h when the drug is no longer detectable (Gould et al., 2019). The effects of two other putative rapid-acting antidepressant drugs (scopolamine, psilocybin) evaluated in treatment-resistant depression (TRD) patients agree with such a dual mechanism of action (Carhart-Harris et al., 2016; Voleti et al., 2013). The initiation mechanism of *(R,S)-*ketamine/HNK (Li et al., 2010) consists of several early synaptic plasticity changes that would contribute to a strengthening of mood-related synapses in the mPFC, then drive the sustained (at t24h) antidepressant actions (expression mechanism). Accordingly, 24, 48 or 72 hours after ketamine administration, there are no measurable levels or trace levels of ketamine or its metabolites in mice, while the antidepressant-like behavioral actions are still present (Gould et al., 2019; Nguyen et al., 2023; Pham et al., 2018). Thus, from this perspective, tissue distribution and clearance parameters quantified during the first 4 hours post-ketamine administration show a reduced synthesis of HNKs in the presence of CYPI. Thus, we can postulate that HNKs are critical for the induction of synaptic plasticity changes following ketamine administration and the elimination of these metabolites here led to a decrease in behavioral antidepressant and biochemical reduction in ketamine-induced increase in cortical GABA release, which may function as an expression mechanism. We previously found that ketamine itself can increase extracellular levels of both glutamate and GABA at t24h in the mPFC (Nguyen et al., 2023; Pham et al., 2018). Here, fluconazole prevented ketamine-induced GABA, but not glutamate, release, and blocked the sustained antidepressant-like effects of ketamine at t24h. Many molecular and cellular changes occur in the time window from 0 to 24 hours post-dose that can explain our data. These results suggest that increases in mPFC GABAergic signaling trigger the sustained effects of ketamine via its (*2R,6R*)-HNK metabolite.

These findings agree with dysregulation of GABAergic and glutamatergic neurotransmission described in several psychiatric disorders, including major depressive episodes (Moriguchi et al., 2019; Sharmin et al., 2023; Yan and Rein, 2022).

In conclusion, we report that (*6*)-HNKs metabolites via an enhanced extracellular GABA levels in the mPFC, exert a significant contribution to the ketamine’s sustained antidepressant activity. Nonetheless, further studies are necessary to elucidate mPFC network organization and its connections with other brain regions to modulate synergistic changes of the balance of excitatory/inhibitory leading to the sustained antidepressant effects of ketamine. The present study also suggests that the dosage of ketamine should be reduced in MDD patients receiving CYPI such as fluconazole and other anti-fungal azole drugs. Potential interactions between (*R,S*)-ketamine and pharmacological CYPIs such as fluvoxamine, a CYPI 2C19; fluoxetine, paroxetine and duloxetine, CYPI 2D6, may occur in MDD and TRD patients. This remark applies particularly to patients receiving esketamine nasal spray as an antidepressant drug treatment, which must be prescribed with an oral conventional antidepressant treatment such as SSRI/SNRI.

## DECLARATION OF INTEREST STATEMENT

DJD serves as a consultant for Lundbeck, Roche, and Servier. AMG serves as a consultant for Lundbeck and Servier. JM, IM-D, DJD, AMG, and CAD are named on provisional patent applications for the prophylactic use of ketamine and other compounds against stress-related psychiatric disorders. PZ, JH, RM, and TDG are co-inventors in patents and/or patent applications related to the pharmacology and synthesis, crystal structure and use of HNKs and related molecules for the treatment of addiction, depression, anxiety, anhedonia, suicidal ideation and post-traumatic stress disorders.

## Abbreviations

aCSF: artificial cerebrospinal fluid;
AD: antidepressant;
AUC: area under the curve;
AUC_0-2hr_: plasma levels from 0 to 2 hours post-administration
AUC_0-3hr_: plasma levels from 0 to 3 hours post-administration
BBB: blood-brain barrier;
BDNF: brain-derived neurotrophic factor;
C_max_: maximum plasma concentration
CYP: cytochrome P450 enzyme
CYPI: cytochrome P450 enzyme inhibitor;
((*6*)-HNK): (*2S,6S;2R,6R*)-HNK
DRN: dorsal raphe nucleus
FST: forced swim test;
Flc: fluconazole;
F10: Flc10, fluconazole 10 mg/kg
F20: Flc20, fluconazole 20 mg/kg
GABA: gamma-aminobutyric acid;
GABA A-R α5: GABA R receptor, α5 sub-unit
Glu: glutamate;
Gln: glutamine;
HNK: hydroxynorketamine;
(6)-HNKs: (*2R,6R;2S,6S*)-HNK
HPLC: high-performance liquid chromatography;
i.p.: intraperitoneal;
(*R,S*)-ketamine: Ket ketamine;
LLOQ: lower limit of quantification;
LOD: limit of detection;
LOQ: limit of quantification
LPS: lipopolysaccharide;
MDD: major depressive disorder
TRD mPFC: medial prefrontal cortex;
mTOR: mammalian target of rapamycin;
NMDA-R: glutamatergic NMDA receptor;
NMDR-2A/2B: glutamatergic NMDA receptor subunit 2A/2B;
norKET: norketamine;
OF: open field;
PK/PD: pharmacokinetic/pharmacodynamic;
S.E.M: standard error of the mean;
SSRI: selective serotonin reuptake inhibitor;
SNRI: serotonin norepinephrine reuptake inhibitor
t_1/2_: plasma elimination half-life;
T_max_: time maximum plasma concentration;
TRD: treatment-resistant depression;
Veh: vehicle;

## ACKNOWLEDGMENTS

The authors would like to thank Dr. Thu Ha Pham for her advice, and colleagues from the animal care facility of SFR-UMRS “*Institut Paris Saclay d’Innovation Thérapeutique*” of University Paris-Sud for their technical assistance. A special thank you to Audrey Solgadi and Pierre Chaminade from the platform “Service d’Analyse des Médicaments et Métabolites (*SAMM*) “ for setting up the mass spectrometry equipment. Our team CESP/UMR-S 1178, Inserm, Univ Paris-Sud/Paris-Saclay provided the necessary resources to perform this study. Thank you also to François Coudoré for his contribution in the lab. A portion of this work was supported by NIH MH107615 and VA Merit 1I01BX004062 to TDG. The RM laboratory is supported by the NIH Intramural Research Program. The contents do not represent the views of the U.S. Department of Veterans Affairs or the United States Government.

## AUTHORS’ CONTRIBUTIONS

Thi Mai Loan Nguyen, Indira Mendez-David, Jean-Philippe Guilloux, Denis J David, Jaclyn N. Highland, Panos Zanos, Todd D. Gould, Christine Ann Denny and Alain M. Gardier contributed to the conception and design of the study; Thi Mai Loan Nguyen, Josephine Cecilia McGowan, Indira Mendez-David, Céline Defaix, Denis J. David, Jacqueline Lovett, Ruin Moaddel, Jaclyn N. Highland, and Panos Zanos contributed to the acquisition of data. Thi Mai Loan Nguyen and Alain M. Gardier wrote the manuscript. All the authors contributed to analysis of data and reviewing the document for intellectual content.

## ADDITIONAL INFORMATION

Supplementary data includes Table S1-Statistics,Table S2-basal dialysate levels, Figure S1-PK 0-4h and FST in CD-1 mice

## FUNDING RESSOURCES

T.M.L.N. was supported by a fellowship from the “France-Vietnam Excellence Scholarships Program”. C.D., I.M-D., L.T., I.E., J-C.A., E.C., D.J.D., A.M.G. from MOODS team, UMR 1018, CESP were supported by Inserm and Université Paris-Saclay to perform this study. This work was also supported by NIH/NIMH R01MH107615 and U.S. Department of Veterans Affairs Merit Awards 1I01BX004062 and 1I01BX006018 to TDG. The contents of this manuscript do not represent the views of the U.S. Department of Veterans Affairs or the United States Government. TDG and PZ are listed as an inventor in patents and patent applications related to the pharmacology and use of a ketamine metabolite, (*2R,6R*)-hydroxynorketamine, in the treatment of depression, anxiety, anhedonia, suicidal ideation, and post-traumatic stress disorders. They assigned patent rights to the University of Maryland, Baltimore, but will share a percentage of any royalties that may be received by the University of Maryland, Baltimore.

## Supplemental

Table S1 Statistics

Table S2 Microdialysis: Basal values of neurotransmitters in the mPFC at t24h

Fig. S1 Pharmacokinetic profile from 0 to 4h post-dose and FST at t24h in CD-1 mice

## Notes

### Competing Interest Statement

The authors have declared no competing interest.

